# Ubiquitin ligase component LRS1 and transcription factor CrHy5 act as a light switch for photoprotection in *Chlamydomonas*

**DOI:** 10.1101/2020.02.10.942334

**Authors:** Nina Lämmermann, Donat Wulf, Kwang Suk Chang, Julian Wichmann, Junhwan Jang, EonSeon Jin, Andrea Bräutigam, Lutz Wobbe, Olaf Kruse

## Abstract

Survival under excess light conditions requires the light-induced accumulation of protein LHCSR3 and other photoprotection factors, to enable efficient energy-dependent quenching in the green microalga *Chlamydomonas reinhardtii*. Here, we demonstrate that the high light-tolerant phenotype of mutant *hit1* is caused by a de-repression of promoters belonging to photoprotection genes, which in turn results from an inactivation of the E3 ubiquitin ligase substrate adaptor LRS1. Transcriptome analyses of *hit1* revealed massive alterations of gene expression modulation as a consequence of perturbed LRS1 function, indicating its role as a crown regulator. In conjunction with random forest-based network modeling, these transcriptome analyses predicted that LRS1 controls photoprotection gene expression via an algal HY5 homolog as its prime transcription factor target. CrHY5 binds to T-box elements present in the promoters of these genes and its inactivation in the *hit1* mutant via CRISPR-Cas9 genome editing, confirmed the regulatory connection between LRS1 and CrHY5, predicted by the network analysis.

## INTRODUCTION

Light provides the energy driving photosynthesis and it is an important cue controlling many physiological processes in phototrophic organisms. In the seed plant *Arabidopsis thaliana* the E3 ubiquitin ligase substrate adaptor COP1/SPA is a master switch of light signaling, by repressing the accumulation of various transcription factors required for the induction of light-dependent processes like photomorphogenesis or flowering in darkness. Repression is achieved by ubiquitination of target transcription factors and their subsequent proteasomal degradation (Deng et al., 1991; Deng et al., 1992; Chen et al., 2006; Lau and Deng, 2012). Red, blue and UV light perception inactivate the COP1/SPA E3 ligase and relieve repression, by several different mechanisms (Podolec and Ulm, 2018). The bZIP transcription factor HY5 is among the prime targets of COP1-mediated repression and activates photomorphogenesis downstream of phytochrome, cryptochrome and UV-B photoreceptors (Gangappa and Botto, 2016). It is also the downstream effector of other signaling cascades and estimated to directly control ∼3000 genes in *Arabidopsis* (Lee et al., 2007). Among the light-dependent processes vital for microalgal cells is non-photochemical quenching (NPQ), which helps to relieve photosystem excitation pressure under conditions that are sub-optimal for photosynthesis, thereby preventing damage to the photosynthetic apparatus (Wobbe et al., 2016). In the green microalga *Chlamydomonas reinhardtii*, a well-established phototrophic model organism, energy-dependent quenching (qE), an NPQ process essential for survival under excess light conditions, requires LHCSR3 (Peers et al., 2009) and PSBS (Tibiletti et al., 2016; Correa-Galvis et al., 2016) proteins, which accumulate upon exposure to high light (Peers et al., 2009) or UV-B radiation (Tilbrook et al., 2016). High light-induction of LHCSR3 accumulation depends on the blue-light photoreceptor phototropin (Petroutsos et al., 2016), while UV-B induction of PSBS and LHCSR1 proceeds via the UVR8 photoreceptor, probably interacting with Cr-COP1, whose absence from algal cells perturbs UV-B acclimation (Allorent et al., 2016; Tilbrook et al., 2016). Cr-COP1 was originally identified in a high light-tolerant *C. reinhardtii* mutant named *hit1* (<u>hi</u>gh light tolerant 1) as LRS1 (putative Light Response Signaling protein; Phytozome v5.5 identifier Cre02.g085050) (Schierenbeck et al., 2015) and transformation with LRS1 restored UV-B acclimation in the *Arabidopsis* UVR8 mutant (Tilbrook et al., 2016), demonstrating that LRS1 (Cr-COP1) is the microalgal homolog of *Arabidopsis* COP1. The mutant *hit1* carries a point mutation in gene *LRS1*, which causes a proline to arginine exchange within the WD40 domain of protein LRS1 (Schierenbeck et al., 2015). Mutant *hit1* originated from a mutagenesis approach based on UV treatment followed by subsequent mutant selection under high light conditions. Thus, mutant *hit1* displays a high light-tolerant phenotype, explicable by an elevated NPQ capacity (Schierenbeck et al., 2015). In the screen another high light tolerant strain *hit2* was identified, whose genome also contains a point mutation in the gene *LRS1* (Schierenbeck et al., 2015) and which is less susceptible to photoinhibition than a wild-type (Virtanen et al., 2019). A recent study indicated the implication of a CUL4-DDB1^DET1^ E3 ligase complex in the control of LHCSR and PSBS gene expression in *C. reinhardtii*, but the effectors acting downstream of the complex remained elusive (Aihara et al., 2019). In *Arabidopsis* the CUL4-DDB1^DET1^ complex acts as an ubiquitin ligase, but how this complex interacts with the COP1-containing complex (CUL4-DDB1^COP1-SPA1^) is unknown (Dong et al., 2015). Two other recent studies demonstrated that the high light-induced accumulation of PSBS and LHCSR proteins requires a homolog of the *A. thaliana* transcription factor CONSTANS, negatively regulated by a CrCOP1-SPA1 E3-ubiquitin ligase and activated by UV-B perception via UVR8 (Gabilly et al., 2019; Tokutsu et al., 2019). However, work from Allorent *et al* (Allorent et al., 2016) demonstrated that LHCSR3 accumulation under high light does not require UVR8, whereas a clear dependence on PHOT was demonstrated by Petroutsos *et al* (Petroutsos et al., 2016). It was thus suggested that blue light perception via phototropin controls LHCSR3 accumulation, while LHCSR1 and PSBS are regulated via UVR-8 (Allorent et al., 2016). In contrast to previous studies, which identified signaling components via forward genetics screens, we took a distinct approach based on regulatory network construction and transcriptomics followed by reverse genetics. Overall, this approach led to the identification of a novel key mechanism of photoprotection control, repressing LHCSR3 transcription in the absence of light.

## RESULTS

### Constitutive over-accumulation of photoprotective proteins in the *hit1* mutant results in high capacity energy-dependent quenching under stress-free conditions

In a previous study, a constitutive high non-photochemical quenching (NPQ) capacity of *hit* strains could be demonstrated as an explanation for their extreme high light tolerance (Schierenbeck et al., 2015). To get further insights into the type of NPQ altered in mutant *hit1*, NPQ induction was recorded for the mutant and the wild-type strain (Fig. 1a), following photoautotrophic growth at moderate high light (400 µmol photons m^-2^·s^-1^). The mutant showed an elevated quenching capacity already within the first seconds of the measurement, which rapidly relaxed in darkness (Fig. 1a; red circles). This indicated that the main NPQ type affected in *hit1* is energy-dependent quenching qE, which is activated within a few seconds and relaxes very quickly in the absence of light (Wobbe et al., 2016). In the green microalga *C. reinhardtii*, efficient energy-dependent quenching requires the proteins LHCSR3 (Peers et al., 2009) and LHCSR1 (Dinc et al., 2016) besides PSBS (Correa-Galvis et al., 2016; Tibiletti et al., 2016), whose accumulation is induced by light stress (Peers et al., 2009; Maruyama et al., 2014). In addition to light, the supply of inorganic and organic carbon regulates the accumulation of LHCSR proteins, with LHCSR3 light induction being repressed by high inorganic/organic carbon supply, whereas LHCSR1 accumulation is unaffected (Polukhina et al., 2016). Similar to LHCSR3, PSBS accumulation is repressed when cells are exposed to high concentrations of carbon dioxide (Correa-Galvis et al., 2016). Therefore, cultivation of a *C. reinhardtii* wild-type at low light intensities (Fig. 1b; LL; PCS) and with the simultaneous presence of acetate in the growth medium prevented the accumulation of LHCSR protein, while LHCSR accumulated in mutant *hit1*, grown under identical conditions. An increase in light intensity (Fig. 1b; MHL) used for cultivation was accompanied by an accumulation of LHCSR in the parental control strain (PCS). The extent of LHCSR accumulation in *hit1* under these conditions was, however, much higher. Similarly, LHCSR1 accumulated to higher levels in *hit1* vs. wild-type (Fig. 1b; αLHCSR1; MHL). In these conditions, PSBS could be immunodetected in samples of *hit1*, but not in those of the wild-type (αPSBS). Exposure to darkness abolished LHCSR accumulation in the parental control strain but not in mutant *hit1* (Fig. 1b; dark), while PSBS could not be detected in either of the two cell lines. To analyze, whether the accumulation of photoprotective proteins in *hit1* cells is due to an increased level of the respective transcript, RT-Q-PCR analyses were conducted (Fig. 1c). The mRNA of genes *LHCSR3.1/3.2* accumulated to much higher levels in *hit1* compared to its parental strain (84.7 ± 7.9 in *hit1* vs. parental strain set to 1) and a similar trend could be observed for the mRNA encoding PSBS1 (Fig 3b; 994 ± 200 vs. 1 in parental strain). It can therefore be concluded that essential components of the qE mechanism, active in *C. reinhardtii*, are constitutively expressed in the mutant *hit1* (Fig. 1b and c). When moderate light stress is applied, LHCSR proteins accumulate in *hit1* to much higher levels than in the parental strain (Fig. 1b and c). COP1, the higher plant homolog of LRS1 is known to be part of a Cullin-based ubiquitin E3 ligase (CUL4-DDB1^COP1-SPA^) complex, repressing the transcription of various genes, whose encoded proteins are only required in the light. Given that the role of COP1 within the signaling mechanisms underlying ultraviolet light acclimation is highly conserved between the higher plant *A. thaliana* and the green microalga *C. reinhardtii* (Schierenbeck et al., 2015; Tilbrook et al., 2016; Aihara et al., 2019; Gabilly et al., 2019; Tokutsu et al., 2019), it seems reasonable to assume that LRS1 functions as a repressor of light-induced genes in dark-acclimated *C. reinhardtii* cells.

**Fig. 1.**
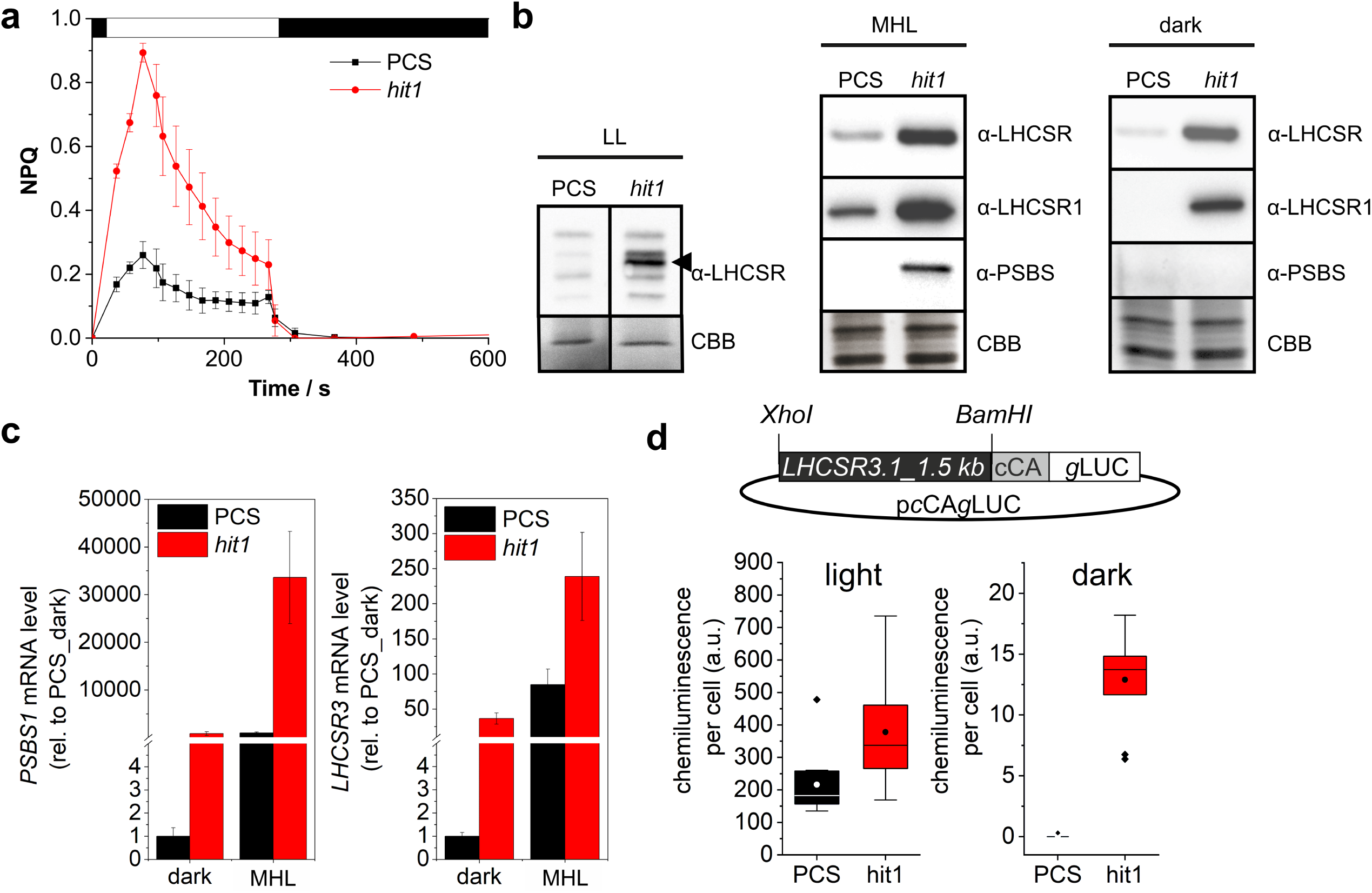
Constitutive over-accumulation of photoprotective proteins in the *hit1* mutant results in rapid activation of non-photochemical quenching. **a**, NPQ induction in phototrophically-grown (400 pmol photons m^2^ s^1^ continuous light) parental control strain (PCS; wild-type *cc124;* black curve) and mutant *hit1* (red curve). NPQ (mean ± standard error; n = 3 biological replicates) was determined during a 5 minutes illumination period (white bar; 1000 pmol m^2^ s^1^ actinic light; 2500 pmol m ^2^ s^1^ saturating pulses) and relaxation monitored by dark incubation (black bar), **b,** Immunodetection of LHCSR1/3 proteins (α-LHCSR), LHCSR1 (α-LHCSR1) and PSBS (α-PSBS) in whole cell protein extracts of PCS and *hit1* grown at either low (LL; 100 pmol photons m^2^ s’^1^) or moderate high light (MHL; 400 pmol photons m^2^ s’^1^) until mid-log phase prior to an 8 hours incubation in darkness. A Coomassie Brilliant blue (CBB) stain served as a loading control. The experiment was repeated three times using distinct biological samples. A representative data set is shown, **c,** Relative *PSBS1* (left panel) and *LHCSR3* (right panel) mRNA levels in PCS and *hit1* (dark-acclimated PCS set to 1; n = 2 biological replicates; mean ± s.d.; Rvalues: *LHCSR3wtvs. hit1* (dark) = 0.003; *LHCSR3* wt vs. *hit1* (dark) = 0.005; *PSBS* wt vs. *hit1* (dark) = 0.01; *PSBS* wt vs. *hit1* (light) = 0.004). Cultures, acclimated to darkness for 8 hours (dark) were exposed to MHL for 0.5 hours, **d,** Upper panel: Reporter construct transformed into *hit1* and its parental strain. Expression of *Gaussia* luciferase *(gLuc)* fused to a carbonic anhydrase secretion signal (cCA) is driven by a 1500 bp sequence located directly upstream of the *LHCSR3.1* start codon. Lower panels: Box-and-whisker-plot showing the interquartile range of the chemiluminescence per cell (mean values of three technical replicates) measured in the supernatant of 10 PCS (black)- or *hit1* (red)-derived transformants after a 5 hours acclimation to moderate high light (light; left panel) or darkness for 72 hours (dark; right panel). The mean chemiluminescence per cell (circles) is shown along with the median (horizontal line) and outliers (diamonds). Rvalues: 0.02 light; 3.92 × 10’^9^ dark.

We therefore reasoned, that a perturbed repressor function of LRS1 in *hit1* might cause a constitutive activation of the *LHCSR3.1* promoter, which is normally only induced by exposure to elevated light or nutrient deprivation (Peers et al., 2009; Grewe et al., 2014; Maruyama et al., 2014). To test this hypothesis, a 1500 bp long region upstream of the start codon of gene *LHCSR3.1* was fused to a *Gaussia* luciferase reporter, containing a secretion signal. Reporter constructs (Fig. 1d; upper panel) were transformed into the *hit1* mutant and its parental control strain to analyze differences in regard to promoter activation in both cell lines. For each transformation experiment (parental strain and mutant) ten transformants were obtained, which stably secreted the luciferase reporter into the culture supernatant. These transformants were exposed to moderate high light for four hours after cultivation in the dark and the luciferase activity was determined in culture supernatants (Fig. 1d; lower part; left panel). In terms of median and mean chemiluminescence per cell as well as in regard to the interquartile range, *hit1*-derived strains showed a higher activity of the *LHCSR3.1* promoter. Importantly, only *hit1*-derived strains expressed the *Gaussia* luciferase to detectable levels in darkness (right panel). This demonstrated that the accumulation of *LHCSR3* mRNA in *hit1* in the absence of excess light as a trigger is the result from an activation of the *LHCSR3* promoter, as could be shown at least for the promoter of *LHCSR3.1*.

### CRISPR-mediated removal of WD40 domains from LRS1 create a *hit*-like phenotype

Considered that its higher plant homolog COP1 acts as a repressor of light-inducible genes, a de-repression of photoprotection genes under dark conditions indicates that the *hit1* mutation in LRS1 is a loss-of-function mutation. In *A. thaliana*, *COP1* null mutations are lethal at the seedling stage, while truncations of the COP1 protein, resulting in variants devoid of the WD40 domain (e.g. *cop*1-4), are viable (McNellis et al., 1994). The WD40 domain is the part of COP1 specifically interacting with target transcription factors and thus required to act as a substrate adaptor for the ubiquitin ligase (Osterlund et al., 1999).

To confirm that the *hit1* mutation, located in the WD40 domain, results in a loss of function, the WD40 domain of LRS1 was completely removed via CRISPR-based *knock-in* at two target sites T1 and T4 (Fig. 2a; Supplemental Material, T1/T4 insertion sites).

**Fig. 2.**
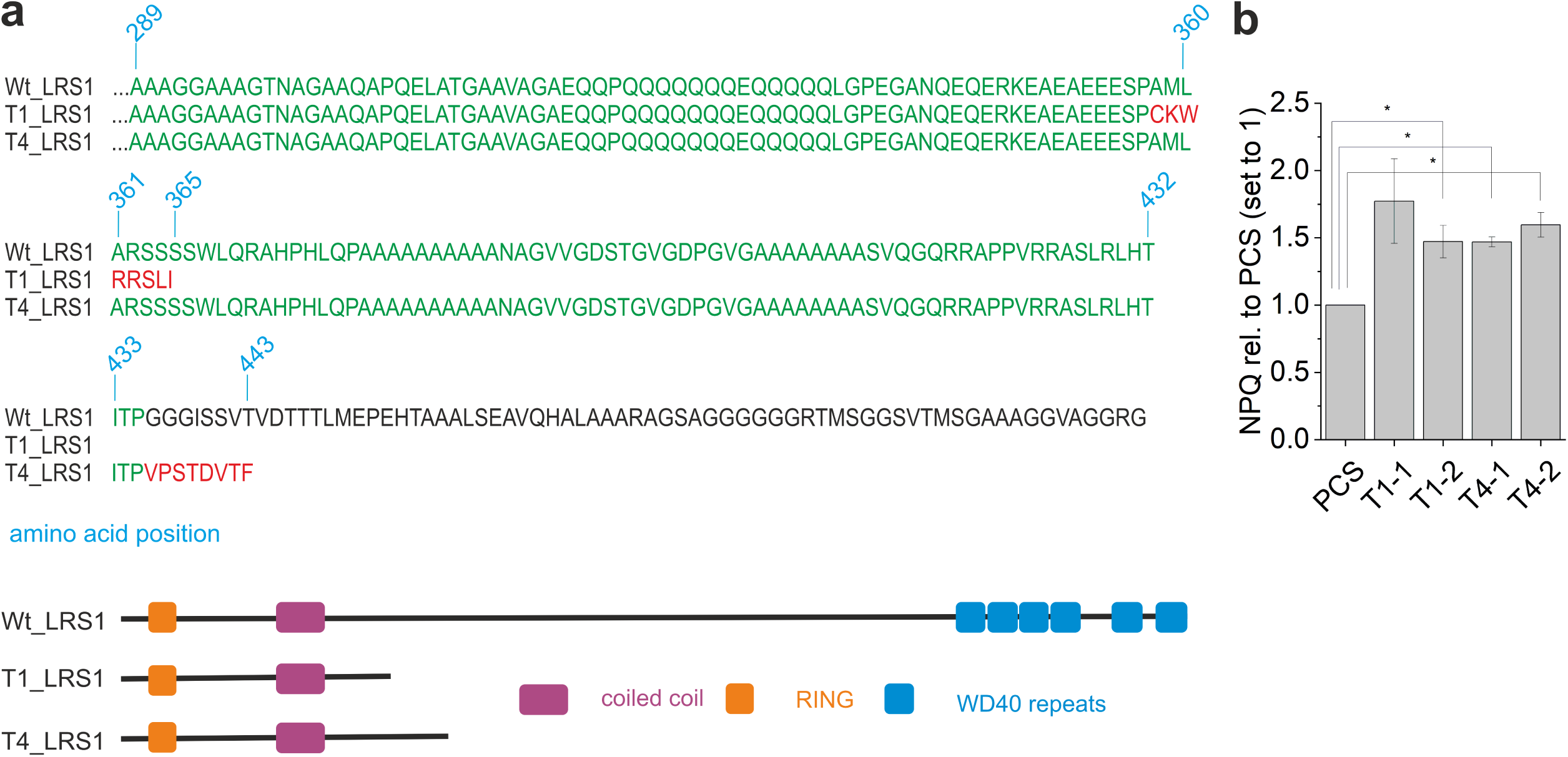
CRISPR-mediated removal of the WD40 domain from LRS1 creates a *hit1*-like phenotype. **a,** Upper part: Amino acid sequence of wild-type (Wt) LRS1 starting a t position 289 and alignment to amino acid sequences created by CRISPR-based gene modification (T1/T4-LRS1) of LRS1 in a wild-type strain. Lower part: Domain composition of Wt_LRS1, T1_LRS1 and T4J.RS1. **b,** Non-photochemical quenching capacity of the parental control strain (PCS) and two selected mutants for each type of gene modification (T1/T4). NPQ of the parental strain was set to 1 and mean values ± s.e. were calculated from three biological replicates. Asterisks indicate significant changes according to a two-tailed Student’s t-test (P<0.05).

As demonstrated by southern blot analysis (Fig. S1), single insertion of a hygromycin resistance cassette at selected target sites resulted in the insertion of a stop codon (T1) or a frameshift mutation (T4) leading to modified *LRS1* genes encoding truncated proteins (Fig. 2a). Truncation of LRS1 in mutants T1 and T4 resulted in a *hit1*-like phenotype with an increased NPQ capacity (≈150%) compared to the parental strain (Fig. 2b), thus demonstrating that the *hit1* mutation located in the WD40 domain (amino acid substitution R→P at amino acid position 1256) (Schierenbeck et al., 2015) interferes with its function.

### A regulatory network for *C. reinhardtii* predicts that photoprotection genes are prime targets of LRS1-mediated repression via transcription factor CrHY5

To investigate a possible effect of the *hit1* mutation on the expression of other genes, we performed a transcriptome-wide analysis based on RNAseq. Mutant *hit1* and the parental control strain were grown in moderate high light (400 µmol m^-2^ s^-1^) before acclimating them to dark conditions for eight hours (t_0_; sampling point “dark”). Cells were then transferred to moderate high light for 45 minutes (t_1_; sampling point “light”) prior to the extraction of RNA. Under dark conditions (t_0_) 1005 genes were up-regulated more than three-fold (log_2_ FC ≥ 1.5; FDR ≤ 0.01) in *hit1* vs. the parental control strain (Supplemental Data Set; “upregulated in *hit1* vs. PCS_dark”). GO Term-enrichment performed with this set of genes indicated a significant enrichment of biological processes such as RNA processing, nucleotide and amino acid metabolism as well as the “generation of precursor metabolites and energy”, which comprises “photosynthesis and light harvesting” as a sub-term. Associated with this GO term are LHCSR proteins, PSBS1, homologs of early light-inducible proteins (ELIPs) (Meyer and Kloppstech, 1984) and the stress-inducible major light-harvesting protein of PSII LHCBM9 (Grewe et al., 2014) (“GO-term enrichment_up_*hit1*_dark”). Among those genes being upregulated in *hit1* under dark conditions were 215 genes which also showed a strong light induction (at least three-fold) in the parental control strain (Supplemental Data Set; “light-induced PCS”). Within this group of genes, however, there was no significant (*P* ≤ 0.01) enrichment of GO terms.

To elucidate the direct targets of LRS1 we employed a machine learning approach. Random forest based regression (Huynh-Thu et al., 2010) predicted interaction partners of LRS1 based on 976 transcriptome samples of wild-type *Chlamydomonas reinhardtii* in public databases (Supplemental Table 1). While COP1-mediated gene repression in *A. thaliana* is known to primarily rely on post-translational processes (Deng et al., 1991; Deng et al., 1992; Chen et al., 2006; Lau and Deng, 2012), we hypothesized that feed-back and feed-forward loops extend to the transcriptome and make it accessible to analysis. The regulatory network predicted by machine learning contained 225941 edges, connecting 17331 nodes. To test whether the network contained relevant biological information, gene enrichment among target genes was tested for each regulatory factor. For 295 out of 812 regulators, enrichment <10^-10^ was detected (Supplemental Table 2) indicating that the predicted network contains biological information. First level, i.e. direct interactors of LRS1 and second level interactors were plotted (Figure 3a; top) to illustrate the network connected to LRS1. 69 target genes directly connect to LRS1 of which 4 are regulators themselves. 620 target genes are indirectly coupled to LRS1 via a connected regulator. The first level interactors were significantly enriched in genes, whose expression was altered significantly with a large amplitude in in the *hit1* mutant (p<10^-30^; Figure 3a, b). The prediction of the network has an average precision for LRS1 of 0.68 for a network cutoff of 0.01 (Figure 3c). CrHY5 is predicted to play a central role in this interaction since it is positioned at the center of interaction from LRS1 to its targets (Figure 3a). The significantly upregulated genes with a large amplitude enrich in the targets of CrHY5 (p<10^-30^). Among the CrHY5 targets predicted by the network are several photoprotection genes (LHCSR3.1, LHCSR1, PSBS2 and other stress-related LHCs; Supplemental Table 3). In addition, other factors required for the induction of energy-dependent quenching via cyclic electron flow such as PGR5 (Johnson et al., 2014) or the synthesis of lutein (FAO5; EC 5.5.1.19 -lycopene beta-cyclase), a xanthophyll implicated in the quenching of excited chlorophyll states (Dall’Osto et al., 2006). Additional differential regulators not captured in the network prediction were pulled from the RNA-seq experiment in *hit1*. These genes and their predicted targets were co-plotted with the LRS1 network (Figure 3a; bottom). RNA-seq data indeed shows that many of their predicted targets are altered in *hit1* as predicted from the network analysis. Network prediction places this sub-network apart from the direct LRS1/CrHY5 nexus. These second level interactors suggested by the RNA-seq data again enrich in de-regulated genes in the network (p<10^-100^). Cr*HY5* (Phytozome ID: Cre06.g310500) is the closest homolog (BLAST analysis; e-value of 8.6·10^-7^) of the *A. thaliana* bZIP transcription factor HY5 (Tair: AT5G11260; UniProt KB O24646; Supplemental Material; Fig. S2), which is a *bona fide* transcription factor target of COP1-mediated photomorphogenesis repression in *Arabidopsis thaliana* (Osterlund et al., 2000).

**Fig. 3.**
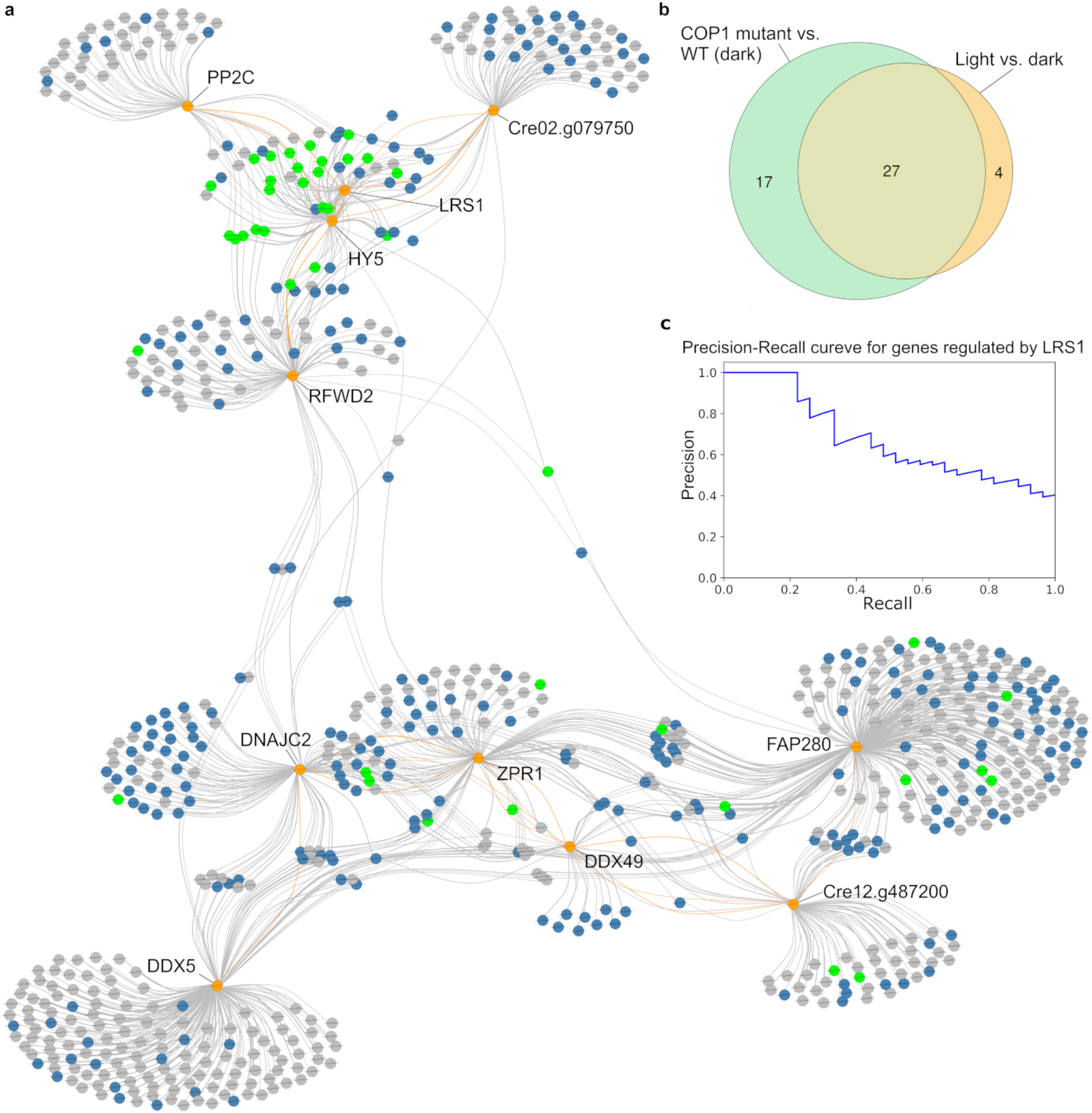
A regulatory network for *C. reinhardtii* predicts that photoprotection genes are prime targets of LRS1-mediated repression via transcription factor CrHY5. **a**, Regulatory network of LRS1 and connected transcription factors (top), sub-network of upregulated regulators (bottom). Interaction of transcription factors (orange) and their targets is shown by a connection between them (green: highly upregulated genes (log_2_>1.5, p<0.01), blue: upregulated genes (1.5>log_2_>0, p<0.01), grey: not differentially expressed), **b,** Venn diagram of significantly upregulated genes (log_2_>1.5, p<0.01) predicted to be controlled by LRS1. Significantly upregulated genes only In the LRS1 mutant vs. the wild-type in the dark (green), significantly upregulated genes in light vs dark (yellow) and the intersection of the differentially expressed genes of the LRS1 mutant and the light dark transition, **c,** Precision-recall curve of genes predicted to be regulated by LRS1. Genes were ranked by their Interaction calculated by genie3. If a gene is significantly upregulated In the mutant and In the light dark transition (intersection from B) it was considered as a true positive. The other predicted interactions were considered as false positives. Precision was calculated cumulatively as the relative amount of true positives.

### The nucleus-targeted bZIP transcription factor CrHY5 is required for the light induction of photoprotection genes in *C. reinhardtii*

The amino acid sequence of CrHY5 contains a putative nuclear localization signal (Fig. S2; red box) and YFP-tagging of CrHY5 in conjunction with confocal fluorescence microscopy confirmed its nuclear localization (Fig. 4a). The transcript of gene Cr*HY5* over-accumulates in dark- and light-grown *hit1* cells compared to its parental strain according to RNAseq results (log2FC *hit1* vs. Wt 2.82 in darkness and 2.81 in the light; Supplementary Data Set). Furthermore, a light induction of Cr*HY5* could be noted for both strains (1.83 wt and 1.81 *hit1*; log_2_FC dark vs. light). To confirm transcript over-accumulation in the mutant, qRT-PCR analyses were conducted with RNA samples derived from dark- and light acclimated cells of *hit1* and Wt (Figure 4b).

**Fig. 4.**
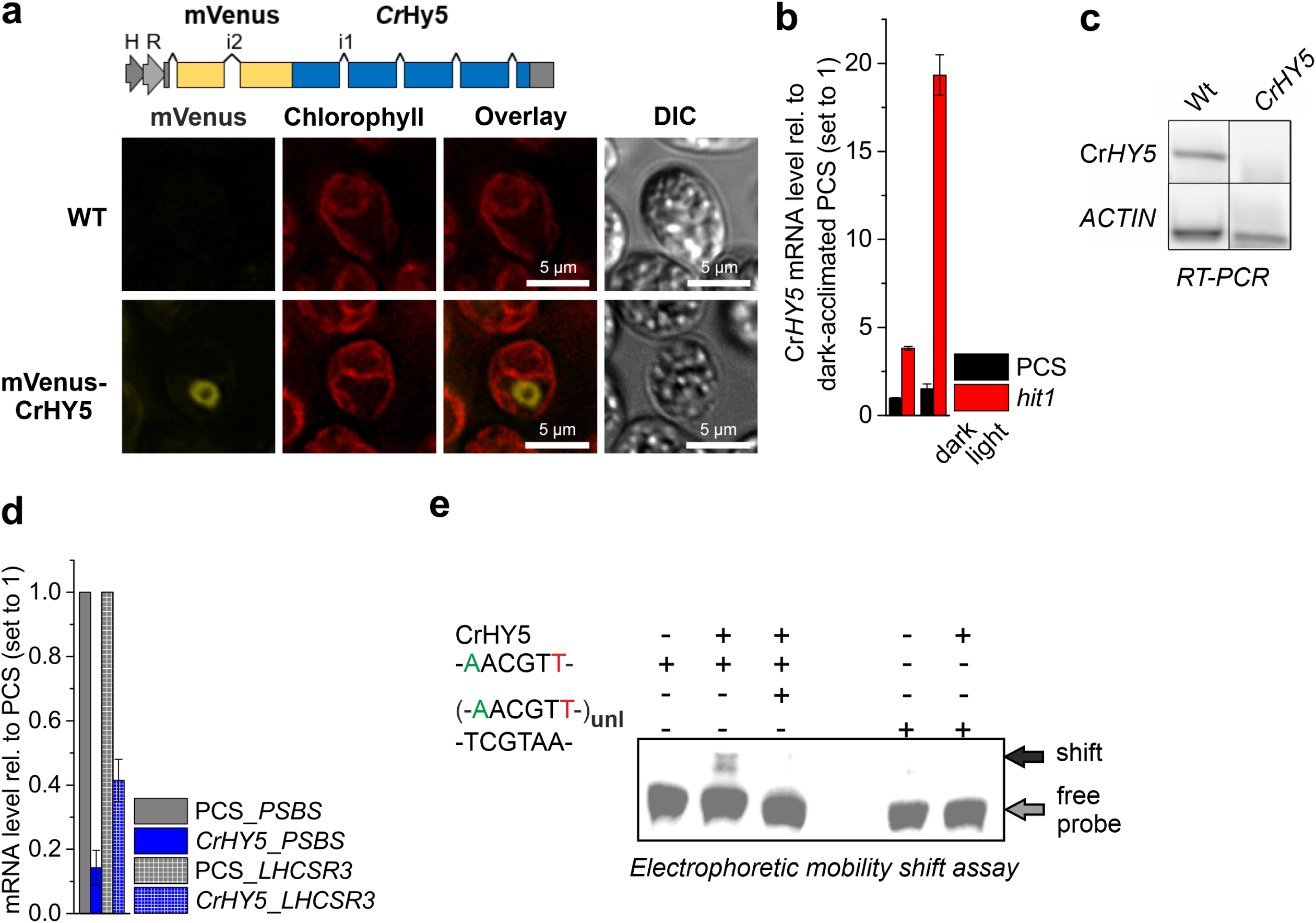
CrHY5, de-regulated in *hit1,* is required for the light induction of photoprotection genes in *Chlamydomonas reinhardtii*. **a**, Scheme depicting the mVenus-CrHY5 construct used for the expression ofthe YFP-tagged protein. Confocal fluorescence microscopy images of a representative individual cell expressing CrHY5 fused to the C-terminus of YFP (mVenus-CrHY5). mVenus fluorescence (mVenus) is shown along with chlorophyll autofluorescence (Chlorophyll), signal overlay (Overlay) and differential interference contrast (DIC). The parental strain (WT) was used as a negative control. Scale bars represent 5 pm. **b,** CrHY5 mRNA accumulation in parental control strain (PCS) and mutant hit 1. mRNA levels are given relative to those determined in dark-grown cultures ofthe parental control strain (set to 1; n = 3 biological replicates; mean ± s.d.). Cultures, acclimated to darkness for 8 hours (dark), were exposed to moderate high light for 0.5 hours, **c,** RT-PCR conducted with primers specific for CrHY5 and total RNAextracted from an insertion mutant (CrHY5; LMJ.RY0402.194448) and its parental strain (Wt). |3-ACTIN served as a housekeeping gene and PCR products were analyzed by agarose gel electrophoresis and DNAstaining after 40 PCR cycles. Representative gel staining from three repeats with distinct samples is shown, **d,** Light-induced mRNA accumulation of PSBS and LHCSR3 determined via RT-qPCR in parental strain (PCS) and mutant CrHY5. Cultures, acclimated to darkness for 8 hours, were exposed to moderate high light for 0.5 hours. mRNA levels (mean values ± s.d. derived from two biological replicates) in PCS were set to 1. **e,** Electrophoretic mobility shift assay with the bZIP domain from protein CrHY5 (25 pmol) and a 26 bp probe (20 fmol) derived from the upstream region of gene LHCSR3.1 containing a T-box element (-AACGTT-) flanked by 10 nucleotides on either side. A 5’-biotin-labelled probe for immunodetection was used along with an unlabeled version (uni.). As a negative control, a mutated version (-TCGTAA-) ofthe probe, lacking the T-box element was tested as well. This position of DNA-protein complexes (shift) and free probe are indicated. A representative shift assay from three individual repeats is shown.

The Cr*HY5* mRNA level was 3.8 (± 0.1; s.d.)-fold higher in dark-acclimated and 19.3 (± 1.1)-fold higher in light-acclimated *hit1* cells, when compared to the wild-type. For HY5 from *A. thaliana*, an autoregulation via binding to its own promoter was shown (Abbas et al., 2014; Binkert et al., 2014). Over-accumulation of CrHY5 in the LRS1 mutant in analogy (Osterlund et al., 2000) to the *A. thalinana* COP1 mutant, might thus result in higher CrHY5 transcript amounts.

Within the first hundred weight-ranked CrHY5 targets predicted by the network several stress-induced light harvesting proteins (LHCSR1, LHCSR3.1, PSBS2, ELI1/3/4/6, HLIP and OHP) could be identified (Supplemental Table 3). To confirm that CrHY5 is indeed required for the light induction of photoprotective genes, a *CrHY5* mutant with a plasmid insertion mapped to the 5’UTR-coding part of the Cr*HY5* gene was used (LMJ.RY0402.194448) (Li et al., 2019). The mutant does not accumulate Cr*HY5* mRNA to levels detectable by RT-PCR (Fig. 4c) and shows diminished levels of *PSBS* (0.14 ± 0.05 s.d. with Wt set to 1) and *LHCSR3* (0.41 ± 0.06) transcripts (Figure 4d). The *A. thaliana* HY5 protein regulates the transcription of light-induced genes such *RIBULOSE BISPHOSPHATE CARBOXYLASE SMALL CHAIN 1A (RBCS1A)* or *CHALCONE SYNTHASE (CHS)* by binding to the G-box light responsive element (LRE) CACGTG (Ang et al., 1998; Chattopadhyay et al., 1998; Lee et al., 2007), which comprises the ACGT core motif recognized by bZIP transcription factors (Foster et al., 1994). Besides the G-box, HY5 was shown to bind other hexanucleotide motifs containing the ACGT core, such as G/T hybrid and C/G hybrid elements (Gangappa and Botto, 2016). Since the whole set of photoprotective light-harvesting genes is up-regulated in mutant *hit1* and these genes are among the CrHY5 targets predicted by the regulatory network, we inspected their 1.5 kb upstream region for the presence of bZIP transcription factor binding sites containing the ACGT core motif (Table S1 and Figure S3). The three most frequent motifs found in the analyzed group of genes were C/G hybrid (12 occurrences), G/T hybrid (8) and G/A hybrid (8) elements (Table S1). The palindromic T-box element AACGTT was exclusively found in the upstream regions of genes *LHCSR3.1* and *LHCSR3.2* (Table S1). Electrophoretic mobility shift assays conducted with the bZIP domain of CrHY5 fused to a GST-tag (Figure 4e) clearly showed that this T-box element is specifically bound by CrHY5.

An impaired light induction of photoprotection genes in the *CrHY5* mutant (Figure 4d), *in vitro* binding to T-box elements present in the promoter regions of genes *LHCSR3.1/2* (Figure 4e) and CrHY5 target predictions from the constructed regulatory network (Figure 3a; Supplemental Table 3) strongly indicate that CrHY5 acts as a transcriptional activator of photoprotection genes. Furthermore, CrHY5 is predicted to be a prime target of LRS1 based on the network analysis (Figure 3a), which is supported by a strong up-regulation of CrHY5 in *hit1* (Figure 4b).

### A knock-out of CrHY5 in the background of a LRS1 truncation mutant abolishes the *hit*-phenotype

To confirm that CrHY5 is indeed a target of LRS1-mediated dark repression and that its accumulation in *hit1* contributes to the over-accumulation of photoprotective factors, we took a genetic approach. To confirm that CrHY5 is a target of LRS1-mediated regulation, we selected the LRS1 truncation mutant *T1-1* to inactivate *CrHY5* in this genetic background. Gene inactivation was conducted via CRISPR-mediated knock-in of a paromomycin cassette into the first exon of gene *CrHY5* (Fig. 5a). Integration of the paromomycin cassette at the desired site was confirmed for the paromomycin resistant mutant *lrs1(T1-1)-hy5* by genomic PCR (Fig. 5b) and sequencing of PCR products (Supplemental Material). The knock-in resulted in the insertion of a stop codon upstream of the bZIP domain encoding part of the gene, modifying it to encode a truncated version of CrHY5, which is devoid of the DNA-binding bZIP domain (Fig. 5c). Along with the parental strain (*lrs1(T1-1)*) and the wild-type used to generate it, mutant *lrs1(T1-1)-hy5* was cultivated under mixotrophic low light and heterotrophic dark conditions, conditions known not to be associated with LHCSR3 accumulation (Peers et al., 2009; Grewe et al., 2014; Maruyama et al., 2014) (Fig. 5d). As expected from the *hit1*-like phenotype (Fig. 2b), the LRS1 truncation mutant *T1-1* displayed a de-repression of *LHCSR3.1* transcription under the examined conditions, while both, the wild-type and the double mutant did not accumulate *LHCSR3.1* mRNA to detectable levels. Abolished *LHCSR3* transcript accumulation following inactivation of *CrHY5* in the LRS1 truncation mutant confirms that CrHY5 is controlled by LRS1 and required for the light induction of *LHCSR3* expression.

**Fig. 5.**
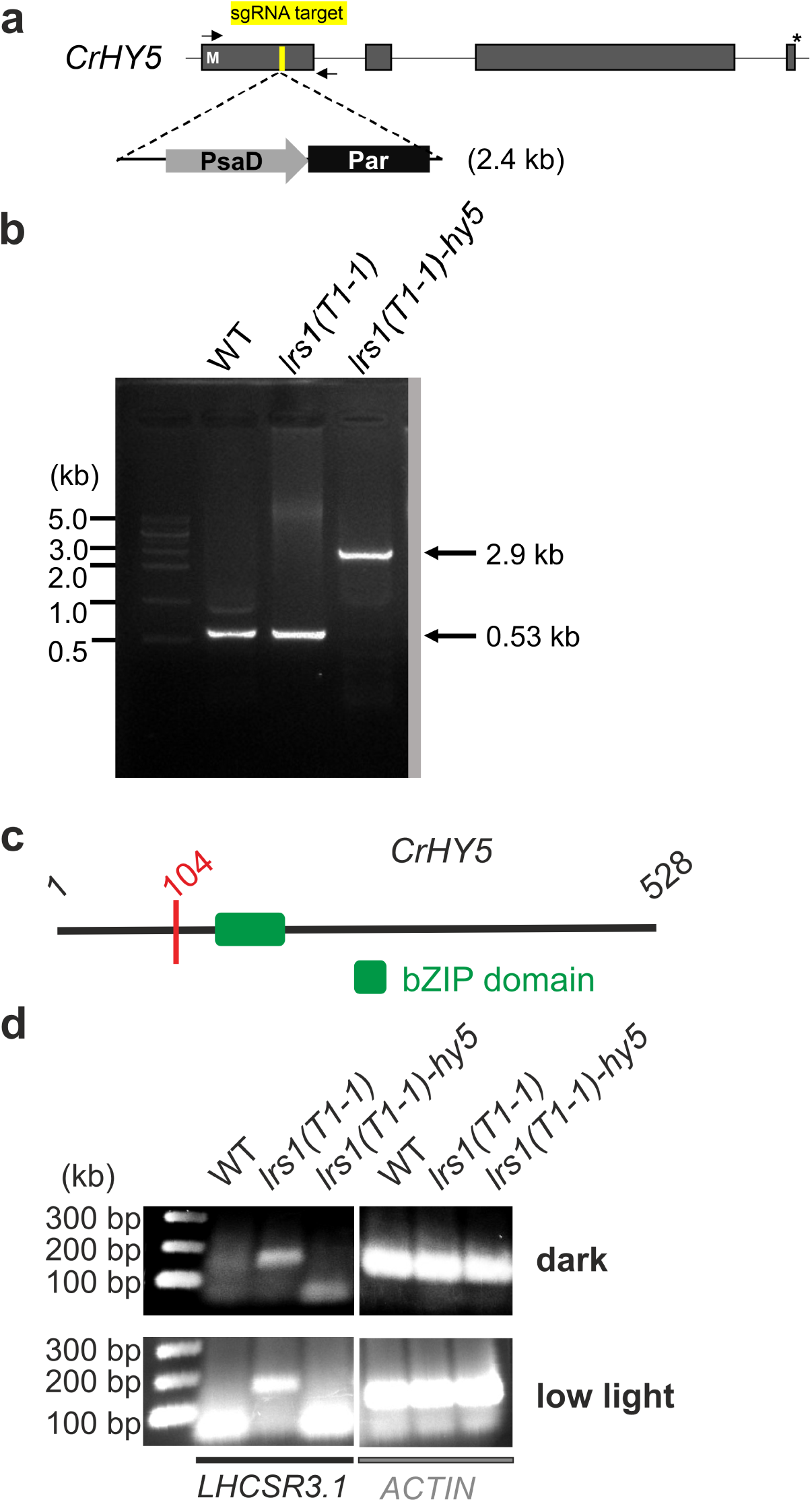
ACRISPR-mediated knock-out of CrHY5 in the background of the LRS1 truncation mutant T1-1 abolishes transcription of *LHCSR3.1* under stress-free conditions. **a**, Schematic representation of the site (sgRNA target; yellow) within the first exon of the *CrHY5* targeted for a knock-in using a paromomycin resistance cassette whose expression is driven by the constitutive *PsaD* promoter. The primers used to obtain the 2.9 kb PCR product after integration of the paromomycin resistance cassette into *CrHY5* are indicated by black arrows, **b,** Results of the PCR performed to confirm the successful integration of the paromomycin resistance cassette (2.9 kb) within the first exon (0.53 kb) of gene *CrHY5.* **c,** Schematic representation of the premature stop codon, which results in the truncation of CrHY5 and subsequent a loss of the bZIP domain, **d,** RT-PCR conducted with primers specific for *LHCSR3.1* and total RNA extracted from mutants T1-1 *(Irs1-(T1-1)),* T1-1 with knock-in of paromomycin resistance cassette in *CrHY5 (Irs1 -(T1-1)-hy5)* and their parental strain (WT). Strains were either incubated for 8 h under dark conditions or cultivated under continuous low light (40 µmol · m^-2^-s^-1^) in TAP medium. *6-actin* was used as the housekeeping gene and PCR products were analyzed by agarose gel electrophoresis and DNAstaining after 40 PCR cycles.

## DISCUSSION

Although, recent studies improved the understanding of photoprotection control in *C. reinhardtii* as a model organism for phototrophic lower eukaryotes (Allorent et al., 2016; Petroutsos et al., 2016; Tilbrook et al., 2016; Aihara et al., 2019; Gabilly et al., 2019; Tokutsu et al., 2019), the depiction of signaling pathways contributing to the light control of *LHCSR3* expression, as key factor, still contains significant gaps. Blue light perception via phototropin was shown to be required for the high light accumulation of LHCSR3 (Petroutsos et al., 2016), but transcription factors acting downstream of PHOT and modulating LHCSR3 expression remained elusive. Similarly, work by Aihara *et al* indicated that E3 ubiquitin ligase components are essential for the dark repression of photoprotection genes including LHCSR3 (Aihara et al., 2019), but could not provide information on downstream effectors. Here, we present a novel LRS1-CrHY5 signaling core, which consists of an E3 ligase component and a bZIP transcription factor. Synoptically, results from the present work allow to depict a new light control mechanism for photoprotection in *C. reinhardtii* (Fig. 6).

**Fig. 6.**
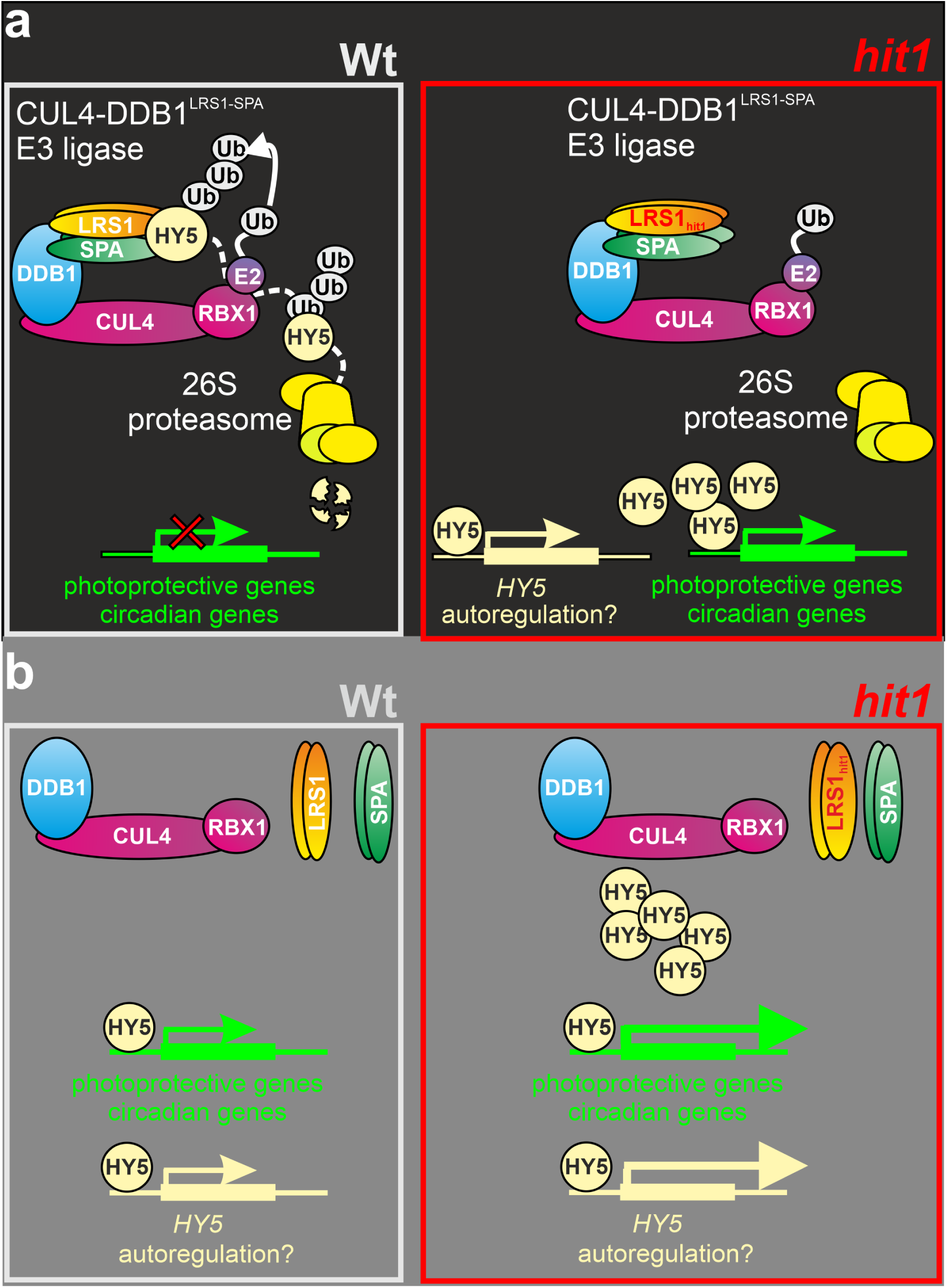
Scheme depicting the repression of photoprotective genes in the nuclei of dark- vs. light-acclimated C. reinhardtii cells by the E3 ubiquitin ligase CUL4-DDB1^LRS1SPA^. **a**, Nuclei of Wt and *hit1* mutant in the dark-acclimated state. Wt: Analogous to the situation on higher plants, a CUL4-DDB1^LRS1SPA^ ubiquitin ligase forms in the nuclei of C. *reinhardtii* and ubiquitinates the transcription factor CrHY5 (HY5), which causes its degradation by the 26S proteasome. This represses the transcription of photoprotection genes, *hitr.* The mutation in LRS1 prevents binding to CrHY5 and thus ubiquitination of the transcription factor. This leads in turn to the accumulation of CrHY5 under dark conditions and transcription of photoprotection genes in the *hit1* mutant. This might also implicate an autoregulation via CrHY5 binding to its own promoter, **b,** Nuclei of Wt and *hit1* mutant in the light-acclimated state. Wt: The CUL4-DDB1^lrs1-spa^ ligase complex dissociates, enabling CrHY5 accumulation and transcription of photoprotective genes. Autoregulation of CrHY5 might enhance induction of photoprotective gene expression, *hitr.h* higher starting amount of CrHY5 after the transfer of dark-acclimated hit1 cells into the light compared to Wt explains the over-accumulation of photoprotective gene transcripts in light-acclimated mutant cells.

Within this mechanism, the E3 ubiquitin ligase component LRS1 and the transcription factor CrHY5 constitute a core interaction and act as a master regulator system for light acclimation in the regulatory network controlling light responses in *Chlamydomonas*. In a wild-type cell (Fig. 6 a; Wt) prolonged darkness should result in the formation of a E3 ubiquitin ligase complex CUL4-DDB1^LRS1-SPA^, whose existence is indicated by the identification of nuclear *Chlamydomonas* genes encoding homologs of the respective complex components known from *A. thaliana* (CUL4**/**Cre12.g516500; DDB1/Cre10.g432000; SPA/Cre13.g602700; RBX1/Cre12.g543450) and the proven interaction of CUL4/DDB1 in Yeast-Two-Hybrid assays (Aihara et al., 2019). The *Chlamydomonas* SPA protein (Cre13.g602700) is among the targets of LRS1 predicted by the network analysis (Supplemental Table 4) and up-regulated in *hit1* (>2-fold under both conditions). A knock-out of SPA was recently shown to de-regulate photoprotection gene expression (Gabilly et al., 2019), confirming the regulatory role of the LRS1/SPA complex for the control of qE. In the same study, an inactivation of *CUL4* (Cre12.g516500) was shown to provoke a similar phenotype. Therefore, the microalgal E3 ligase complex should ubiquitinate transcription factors in analogy to its function in *Arabidopsis* (Deng et al., 1991; Deng et al., 1992; Chen et al., 2006; Lau and Deng, 2012) and with LRS1/SPA acting as a substrate adaptor.

The bZIP transcription factor CrHY5 is predicted to be a prime target of LRS1 (Figure 3), strongly up-regulated in *hit1* (Fig. 4b), required for proper light induction of photoprotection genes (Fig. 4c and d) and essential for constitutive LHCSR3 expression in the LRS1 mutant (Fig. 5d). Thus, we conclude that this transcription factor is a prime regulatory target of the CUL4-DDB1^LRS1-SPA^ complex and dark-repressed via ubiquitination and subsequent proteasomal degradation (Fig 6a, Wt). The point mutation present in the WD40 domain of LRS1 in the *hit1* mutant creates a loss of function by abolishing transcription factor binding (Fig. 6a, *hit1*), which could be confirmed by CRISPR-mediated removal of the WD40 domain region from LRS1, resulting in a *hit1*-like phenotype (Fig. 2b). As a consequence, the impaired function of the CUL4-DDB1^LRS1-SPA^ E3 ligase permits an accumulation of CrHY5 and induction of photoprotection genes under dark conditions. Increased *CrHY5* transcript amounts in *hit1* under dark and light conditions (Fig. 4b) point at an autoregulation similar to the one described for Arabidopsis (Abbas et al., 2014; Binkert et al., 2014). We hypothesize that under excess light conditions the E3 ligase complex dissociates, thus relieving the repression of photoprotection gene expression (Fig. 6b). High light induction of photoprotection genes (Fig. 1b, c, d) and *CrHY5* (Fig. 4b) is amplified in the *hit1* mutant, since the existence of transcription factors preformed via constitutive expression should increase induction strength (Fig. 1b-d). This aspect of the phenotype together with the constitutive presence of LHCSR proteins protects *hit1* from the photoinhibition caused in a wild-type by a sudden change of light intensity from low to very high light, as is tolerated by *hit1* (Schierenbeck et al., 2015).

What is the light signal modulating the activity of the CUL4-DDB1^LRS1-SPA^ complex? LRS1 has been shown to interact with UVR8 upon UV-B exposure (Tilbrook et al., 2016; Tokutsu et al., 2019) and in *Arabidopsis* this interaction is proposed to sequester the substrate adaptor COP1-SPA1 from the ubiquitin ligase complex, which stabilizes HY5 as a key transcription factor governing UV-B acclimation (Huang et al., 2013; Podolec and Ulm, 2018). Interestingly, as HY5 in *A. thaliana*, CrHY5 is strongly induced by UV-B light exposure of *Chlamydomonas* cells (Tilbrook et al., 2016) suggesting a conservation of this signaling mechanism between seed plant and green microalga and that CrHY5 might be vital for UV-B acclimation besides CONSTANS (Tokutsu et al., 2019), which is, however, not among the first-level interactors of LRS1 within the regulatory network (Supplemental Table 4). In contrast to LHCSR1 and PSBS, the accumulation of LHCSR3 is rather induced by excess light of longer wavelengths than UV-B radiation (Allorent et al., 2016) and phototropin-mediated blue light perception was shown to be critical for this regulation (Petroutsos et al., 2016). Phototropins are not known to interact with E3 ligase complex components and PHOT is not among the predicted interactors of LRS1 (Supplemental Table 4), while other light receptors are (e.g. CRY-DASH1/Cre02.g078939). Future work will address the identification of light receptors acting upstream of the LRS1-CrHY5 signaling core, which will be facilitated by knowledge about the topology of the LRS1 signaling network and existing mutants.

## METHODS

### Strains and Growth Conditions

*Chlamydomonas reinhardtii* wild type strain *cc124, Hy5* mutant strain (LMJ.RY0402.194448) (Li et al., 2019) with an insertion in the 5’ UTR of gene Cre06.g310500 and its parental strain cc4533 were obtained from the *Chlamydomonas* resource center. Mutant *hit1* was generated from parental strain *cc124* via UV mutagenesis (Schierenbeck et al., 2015). Cells were either grown photoautotrophically in high salt medium HSM (Harris, 1989) bubbled with 2% (v/v) CO_2_ or mixotrophically in tris acetate phosphate (TAP) medium (Harris, 1989) using the indicated light intensity.

### Chlorophyll and Fluorescence Measurements

Prior to chlorophyll fluorescence measurements, cells suspensions (5 µg Chl/mL) were acclimated to darkness for 10 min. NPQ measurements were performed with Mini-PAM (Walz, Effeltrich, Germany) using a standardized program for recording a dark-to-light induction curve for 5 min followed by a dark recovery phase with the software WinControl (V2.08.). Dark adapted cells supplemented with 10 mM bicarbonate were illuminated with pulsed actinic white light at 1000 µmol m^-2^ s^-1^. During the dark recovery phase consecutive saturated pulses were applied at 10 sec, 30 sec, 1 min, 2 min and 5 min after termination of illumination. NPQ was calculated by the equation (F_m_-F_m’_)/F_m’_ in which F_m_ is the maximal fluorescence yield of dark-adapted cells and F_m’_ is the maximal fluorescence yield recorded during actinic light illumination (Maxwell and Johnson, 2000).

### Protein quantification via immunoblotting

*cc124* and *hit1* cultures were grown mixotrophically in TAP under continuous light (100 µmol m^-2^ s^-1^) and constant agitation (120 rpm) until they reached an OD_750_ of 0.4 (low light sample). They were then transferred into complete darkness for 8 hours (dark sample) and subsequently transferred into moderate high light conditions (400 µmol m^-2^ s^-1^) for 2h (MHL sample). For the preparation of SDS-PAGE samples, cell pellets containing 3 × 10^7^ cells were resuspended in 150 μL lysis buffer (2% (w/v) SDS, 60 mM Tris-HCl, pH 6.8, 100mM dithiothreitol, and 10% (w/v) glycerol) and boiled for 5 min at 95°C. The protein concentration in samples was determined using the DC Protein Assay (BioRad, CA, USA). Protein samples (10 μg per lane) were separated on 10% (w/v) or 15% (w/v) denaturing Tris-glycine SDS-PAGE gels (Laemmli, 1970) stained with Coomassie Brilliant Blue R-250 or electroblotted onto nitrocellulose membranes (0.2-mm pore size; GE Healthcare, Chicago, Illinois). Immunoblotting analyses were performed using specific primary antibodies and a rabbit-specific horseradish peroxidase-conjugated secondary antibody (Agrisera, Vännäas, Sweden). Signals were visualized using the FUSION-FX7 detection system (Vilber Lourmat, Eberhardzell, Germany). Rabbit anti-LHCSR1 (Agrisera; product AS14 2819), rabbit anti-LHCSR (Bonente et al., 2011) and rabbit anti-PSBS (Bonente et al., 2008) were kindly provided by R. Bassi (Verona, Italy).

### RNA isolation for mRNA quantification

*cc124* and *hit1* cultures were grown mixotrophically in TAP under continuous light (100 µmol m^-2^ s^-1^) until they reached an OD_750_ of 0.4 and transferred into darkness for 8 hours (dark sample) before they were exposed to moderate high light conditions (400 µmol m^-2^ s^-1^) for 30 min (light sample). Total RNA was isolated according to Chomczynski and Sacchi (Chomczynski and Sacchi, 2006) with integrated RNase free DNaseI (Promega, Madison, Wisconsin) treatment.

For RTqPCR, 200 ng total RNA was used in each single-step reaction according to the manufacturer’s instruction (HiRoxSensifast Kit, Bioline). Primers (LHCSR3 fw/rev: 5’-ggcaccggtttcaactccct-3’/ 5’-cggccgttgttcagctcctt-3’; PSBS1 fw/rev: 5’-aggcgacccacaaacagctc-3’/ 5’-cgcatcaccgtgcacttcaac-3’; Hy5 fw/rev: 5’-tgagcaagctgagtccggag-3’/ 5’-ttccgcagtgcctcgttctg-3’; β-actin fw/rev: 5’-atgggccagaaggactcgtac-3’/ 5’-tcgttgaagaaggtgtggtg-3’) were designed using Primer3web (http://primer3.ut.ee/) (Kõressaar et al., 2018) or NCBI Primer Blast (https://www.ncbi.nlm.nih.gov/tools/primer-blast/) (Ye et al., 2012). β-Actin was used as a housekeeping gene and relative expression values determined according to Pfaffl *et al*. (Pfaffl, 2001).

### RNA-Seq Experiments

cDNA library and illumina sequencing were performed as previously described (Jaeger et al., 2017b). For each timepoint three biological replicates were used and processed individually. The library pool was sequenced 2x 70 nt paired-end using rapid run mode on Illuminàs HiSeq 1500 50% (Illumina, San Diego, California). Paired end reads of triplicated Wt and *hit1* samples were mapped onto the transcriptome v5.5 of *Chlamydomonas reinhardtii* with kallisto (Bray et al., 2016) (v44; default parameters). Differential expression was determined by edgeR (Robinson et al., 2010) (classic mode; common dispersion followed by tag-wise dispersion) followed by Benjamini-Hochberg multiple hypothesis testing correction (Benjamini and Hochberg, 1995). Enrichment analyses were conducted with topGO (Adrian Alexa, 2017) using Fishers Exact Test (Fisher, 1922). RNAseq data were submitted to the NCBI SRA archive (https://www.ncbi.nlm.nih.gov/bioproject/596622).

### Regulatory network construction

The network was constructed based on publicly available data. 1206 short read archive files (retrieved 10/15/2018) were screened for non-mutant files to avoid network construction errors which resulted in 1050 files for analysis. Reads were mapped on the primary transcriptome v5.5 using kallisto (Bray et al., 2016). The resulting expression matrix was screened in gene direction by selecting only genes which exceeded a tpm of 5 in at least one experiment to include only genes expressed above background. To allow for sufficient read depth per experiment, only experiments with at least 10^6^ mapped reads were kept. Transcription associated proteins required for prediction were pulled from https://plantcode.online.uni-marburg.de/tapscan/ (retrieved 10/19/2018) (Wilhelmsson et al., 2017). 975 experiments with 17731 transcripts were used to predict the regulatory interactions using genie3 (Huynh-Thu et al., 2010) with the transcription associated proteins as input genes, the random forest method (Breiman, 2001) with 1000 trees and the square root of the transcription associated proteins as input genes as regulators for each tree. Interactions above the cut-off of 0.005 were used for analyses. Gene ontology enrichment analysis using the R package topGO (Adrian Alexa, 2017) with the classic algorithm and the fisher statistic was used to functionally annotate all regulatory factors. For differential expression analysis, RNA-seq reads were mapped using the pipeline described above. Differential expression was tested with classic edgeR (Robinson et al., 2010) (counts were normalized using estimateCommonDisp and estimateTagwiseDisp sequentially and tested using exactTest). Venn diagrams were constructed with the R package eulerr.

### Luciferase activity assay

For obtaining a plasmid expressing *Gaussia princeps* luciferase (gLuc) under control of the LHCSR3.1 (Cre08.g367500) promoter (*Pro_LHCSR3.1_:cCAgLuc*) a 1.5-kb element upstream of the LHCSR3.1 translation start codon ATG was amplified from *C. reinhardtii* cc124 genomic DNA with primers containing *XhoI* and *BamHI* restriction sites. For insertion of required restriction sites, vector p*OptgLuc* (Lauersen et al., 2015) was amplified with primers containing *BamH*I and *Bgl*II restriction sites as well as a native *C. reinhardtii* secretion signal (Lauersen et al., 2013). *LHCSR3.1* promoter was then placed upstream of the secretion signal and reporter gene by digesting vector and 1.5-kb element with *BamH*I and *Xba*I FastDigest® restriction endonucleases (Thermo Scientific, Waltham, Massachusetts). Construct *Pro_LHCSR3.1_:cCAgLuc* was then transformed into cc124 and *hit1* by electroporation as previously described (Jaeger et al., 2017a). Positive transformants expressing the luciferase under control of the *LHCSR3.1* promotor were detected using agar plate-based bioluminescence assay (Lauersen et al., 2013). Twelve *Pro_LHCSR3.1_:cCAgLuc* transgenic cc124- and *hit1*-derived transformants, respectively, were grown mixotrophically in 12-well plates until OD_750_ of 0.5 with continuous illumination (100 µmol m^-2^ s^-1^) and constant agitation (150 rpm). Cells were centrifuged (2 min, 2000 × g), supernatant discarded, resuspended in fresh TAP media and partitioned into well-plates. The cells were incubated either for three days in complete darkness or for 4h at 400 µmol m^-2^ s^-1^ with continuous agitation. Cell suspension samples were taken and centrifuged at 10,000 *g* to obtain cell-free supernatants. To 100 µL supernatant 80 µL buffer (0.1 M K_2_HPO_4_ (pH 7.6), 0.5 M NaCl, 1 mM EDTA) and 20 µL coelenterazine solution (0.01 mM) were added. Luminescence was recorded for 10 s in a Fluostar Optima Spectrometer (BMG Labtech, Ortenberg, Germany).

### Electrophoretic mobility shift assays with the bZIP domain of CrHY5

For electro mobility shift assay studies (EMSA) the *E. coli* codon optimized bZip domain of protein Cre06.g310500.t1.2 (amino acid position 141-203) was cloned into the GST-containing vector pGEX-4T-2 (GE Healthcare Life Sciences) by the company GeneScript. *E. coli* strain BL21DE3 served as a host for protein expression at 18°C for 16 h. Protein purification was performed as previously described, using sonication and GST-affinity chromatography, respectively (Wobbe and Nixon, 2013). EMSA studies were performed using the LightShift Chemiluminescent EMSA-Kit (Thermo Scientific) according to the manufacturer’s instructions. For binding reactions 800 ng protein were used with 4 pmol unlabeled probe and 20 fmol biotinylated probe.

### Fluorescent protein-tagging of *Cr*Hy5

The nucleotide sequence of *C. reinhardtii* Hy5 (Uniprot: A0A2K3DRN7) was optimized as previously described (Baier et al., 2018) using the Intronserter tool (Jaeger et al., 2019) (https://bibiserv.cebitec.uni-bielefeld.de/intronserter). The codon optimized, intron containing gene was chemically synthesized (GenScript, Piscataway Township, New Jersey) and cloned between *Bam*HI-*Bg*lII or *Eco*32I-*Eco*RI in the pOpt2_mVenus_Paro vector (Wichmann et al., 2018). *C. reinhardtii* strain UVM4 (Neupert et al., 2009) (graciously provided by Prof. Dr. Ralph Bock) was transformed with glass bead agitation (Kindle, 1990) and positive transformants were identified by fluorescence microscopy screening for the mVenus (YFP) reporter (Lauersen et al., 2016; Wichmann et al., 2018). Localization of the mVenus-linked proteins was confirmed by wide-field fluorescence microscopy (Lauersen et al., 2016).

### Targeted mutagenesis of Chlamydomonas reinhardtii

For the mutagenesis experiments*, Chlamydomonas reinhardtii* cw15 mt^-^ (*cc4349*) was used as the wild type. *C. reinhardtii cc4349* mutants were generated using targeted-DNA knock-in mutagenesis through the direct transformation of the pre-assembled Cas9 protein-sgRNA ribonucleoproteins (RNP) complex and PCR fragments carrying a hygromycin or paromomycin resistance gene. The PCR products using the pChlamy_3 vector (Invitrogen, Thermo Fisher), which contained the expression cassette for the hygromycin (Hyg) resistance gene, were introduced into the parental strain, CC-4349. Specific primers for the Hyg vector (Fw: 5‘-atgattccgctccgtgtaaatg-3’ and Rv: 5‘-agtaccatcaactgacgttacattc-3’) were used for PCR amplification. The PCR product encoding paromomycin resistance gene (Par) was amplified from the pPEARL (GeneBank accession: KU531882) with Par primers (Fw: 5’-gaacacgcaggtcagacc-3’ and Rv: 5’-gtccacactgtgctgtcac-3’). CRISPR-Cas9 driven mutagenesis was performed based on the method in Baek *et al*. 2016 (Baek et al., 2016) and Yu *et al*. 2017 (Yu et al., 2017). To generate target-specific single guide RNAs (sgRNAs) for *CrLRS1* or *CrHY5*, the sgRNAs were designed using Cas-Designer, a web-based CRISPR RGEN tool, (http://www.rgenome.net/cas-designer/) (Park et al., 2015). The selected target sites for the *CrLRS1*-sgRNA-T1 target were as follow: 5’-ggaggagtcgccagctatgctgg-3’, *CrLRS1*-sgRNA-T4 target: 5’-ctgcacacgatcacgccaggtgg-3’, and *CrHY5*-sgRNA target: 5’-tctgaggagtcgtcatcccgcgg-3’. The *CrLRS1*-specific or *CrHY5*-specific sgRNAs were synthesized using the GeneArt™ Precision gRNA Synthesis Kit (Invitrogen, Thermo Fisher) following the manufacturer’s instructions. The Cas9 protein was purchased from ToolGen Inc. (Seoul, South Korea). The Cas9 protein (100 µg) and *in vitro* synthesized sgRNA (40 µg) were incubated for 10 min at room temperature. *C. reinhardtii* (1× 10^7^ cells) were electroporated with RNP complex and PCR fragments as described by Baek *et al*. 2016 (Baek et al., 2016). After transformation, the cells were resuspended in TAP-40 mM sucrose solution were incubated for 16 hours with gentle agitation and transferred onto TAP plate containing antibiotics. For the *CrLRS1* targeted mutants, colonies with *lrs1* are selected on the hygromycin media. However, for the *CrHY5* targeted double knock-out mutant *(lrs1-(T1-1)-hy5)* generated from the *lrs1* background, selections were carried out using paromomycin media. After 5 days of incubation, the surviving colonies were screened for the mutant selection by PCR analysis.

#### Genotypic characterization of mutants

Genomic DNA of *C. reinhardtii* was extracted according to the protocol by Baek *et al*. 2016 (Baek et al., 2016) for genotypic characterization of the wild type and mutants. For detection of the targeted knock-in mutations, the target regions were amplified by PCR with LRS1-specific primers (Fw: 5’-cgagcctggagtacgacatcat-3’, and Rv: 5’-gtgctctggctccatcagtgta-3’), and HY5-specific primers (Fw: 5’-atggactctagcactgctgc-3’ and Rv: 5’-tgactccaagaccagcgaac-3’). The PCR products were sequenced using Sanger sequencing by Macrogen Inc. (Seoul, South Korea). For Southern blot analysis, genomic DNAs (20 µg) of wild-type and *lrs1* mutants digested with *Pst* I and the fragments were analyzed as described by Seo *et al*. 2018 (Seo et al., 2018). The partial hygromycin gene was used as a probe, which was amplified by PCR with primers (Fw: 5’-atgattcctacgcgagcctg-3’ and Rv: 5’-atccggctcatcaccaggta-3’).

## Accession Numbers

*The following Phytozome v5.5 identifiers were used: LRS1/*Cre06.g310500; *CrHY5*/*Cre02.g085050; LHCSR1/Cre08.g365900.t1.2; LHCSR3.1/Cre08.g367500.t1.1; LHCSR3.2/Cre08.g367400.t1.1; PSBS2/Cre01.g016750.t1.2; ELI1/ Cre08.g384650.t1.2; ELI3/Cre09.g394325.t1.1; ELI4/ Cre02.g143550.t1.2; ELI6/ Cre17.g740950.t1.2; HLIP/ Cre02.g109950.t1.2; OHP1/ Cre06.g251150.t1.2*.

RNAseq data were submitted to the NCBI SRA archive (https://www.ncbi.nlm.nih.gov/bioproject/596622).

## SUPPLEMENTAL DATA

**Supplemental Dataset 1.** Summary of expression estimates (tpm) of 13055 genes showing a significantly different expression (q≤0.01) for at least one examined condition (*hit1* vs. PCS in darkness/light; dark-to-light transition for *hit1*/PCS)

**Supplemental Table 1.** List of SRA accessions (https://www.ncbi.nlm.nih.gov/sra) used for network construction

**Supplemental Table 2.** GO term enrichment table

**Supplemental Table 3.** Prime targets of CrHY5 according to their weight score in the network analysis sorted by their assigned function

**Supplemental Table 4.** Prime targets of LRS1 according to their weight score in the network analysis sorted by their assigned function

## Supplemental Material (Word Document)

**Supplemental Table 5:** ACGT motifs present in the region 1.5 kb upstream of transcription start sites of photoprotection genes.

**Supplemental Sequence 1:** T1 insertion site.

**Supplemental Sequence 2:** T4 insertion site.

**Supplemental Sequence 3:** Insertion site of *CrHY5* with Par gene in the *CrLRS1* mutant, *lrs1(T1-1)*.

**Supplemental Figure 1:** Southern blot analysis of wild-type and *CrLRS1* mutants.

**Supplemental Figure 2:** Alignment of CrHY5 and AtHY5

**Supplemental Figure 3:** Localization of ACGT motifs within the promoters of photoprotection genes

## AUTHOR CONTRIBUTIONS

N.L., L.W. and O.K. conceived and designed the research. N.L. performed most of the experiments. D.W. and A.B. constructed the regulatory network and analysed RNAseq data. K.S.C, J.J. and E.J. generated CRISPR mutants and analysed their genotype.

J.W. performed localization experiments. L.W. and O.K. wrote the manuscript, with input from the co-authors.

## ACKNOWLEDGMENTS

We thank R. Bassi (University of Verona) for the provision of antisera. We are grateful to the Center for Biotechnology (CeBiTec) at Bielefeld University for access to the Technology Platforms.

